# Dystrophin regulates peripheral circadian SRF signalling

**DOI:** 10.1101/2021.06.23.449492

**Authors:** Corinne A Betts, Aarti Jagannath, Tirsa LE van Westering, Melissa Bowerman, Subhashis Banerjee, Jinhong Meng, Maria Sofia Falzarano, Lara Cravo, Graham McClorey, Katharina E Meijboom, Amarjit Bhomra, Wooi Fang Lim, Carlo Rinaldi, John R Counsell, Katarzyna Chwalenia, Elizabeth O’Donovan, Amer F Saleh, Michael J Gait, Jennifer E Morgan, Alessandra Ferlini, Russell G Foster, Matthew JA Wood

## Abstract

Dystrophin is a sarcolemmal protein essential for muscle contraction and maintenance, absence of which leads to the devastating muscle wasting disease Duchenne muscular dystrophy (DMD)[1, 2]. Dystrophin has an actin-binding domain [3–5], which specifically binds and stabilises filamentous (F)-actin[6], an integral component of the RhoA-actin-serum response factor (SRF)-pathway[7]. The RhoA-actin-SRF-pathway plays an essential role in circadian signalling whereby the hypothalamic suprachiasmatic nucleus, transmits systemic cues to peripheral tissues, activating SRF and transcription of clock target genes[8, 9]. Given dystrophin binds F-actin and disturbed SRF-signalling disrupts clock entrainment, we hypothesised that dystrophin loss causes circadian deficits. Here we show for the first time alterations in the RhoA-actin-SRF-signalling-pathway, in both dystrophin-deficient myotubes and dystrophic mouse models. Specifically, we demonstrate reduced F/G-actin ratios and nuclear MRTF, dysregulation of core clock and downstream target-genes, and down-regulation of key circadian genes in muscle biopsies from DMD patients harbouring an array of mutations. Further, disrupted circadian locomotor behaviour was observed in dystrophic mice indicative of disrupted SCN signalling, and indeed dystrophin protein was absent in the SCN of dystrophic animals. Dystrophin is thus a critically important component of the RhoA-actin-SRF-pathway and a novel mediator of circadian signalling in peripheral tissues, loss of which leads to circadian dysregulation.

## Introduction

Skeletal muscle is a dynamic structure in which myofilament turnover, maintenance and energy replenishment occur continually throughout the day [10]. Circadian transcriptomic studies in skeletal muscle indicate that approximately 3.4% of expressed skeletal muscle genes show rhythmicity, and that this differs between muscle types, specifically slow and fast muscle [11]. These rhythmic genes are involved in many central processes such as myogenesis, muscle lipid utilisation, protein metabolism and organisation of myofilaments, and very recently a key circadian gene, *Bmal1* (*Arntl*), was shown to be involved in impaired myogenicity in muscle of dystrophic mice [12].

Dystrophin is an integral sarcolemmal protein essential for muscle contraction and maintenance, absence of which leads to the devastating muscle wasting disease Duchenne muscular dystrophy (DMD)[1, 2]. Dystrophin has an actin-binding domain at the amino-terminus of the full-length isoform [3–5], which specifically binds and stabilises filamentous (F)-actin [6], an integral component of the RhoA-actin-serum response factor (SRF)-pathway [7–9]. The RhoA-actin-SRF-pathway is well described in muscle [13], and has since been shown to play an essential role in circadian signalling via systemic cues activating SRF in peripheral tissues [8]. Indeed SRF is a pivotal nuclear transcription factor, regulating over 200 target genes that are predominantly involved in cell-growth, migration, cytoskeletal organisation and myogenesis, and one of the earliest SRF target genes to be identified was *Dmd* [13]. This integral relationship, combined with the understanding that they operate via a feed-back loop, intimates that the absence of dystrophin would have serious implications on SRF regulation. Interestingly, studies designed to mimic age related sarcopenia by disrupting skeletal muscle SRF expression resulted in atrophy, fibrosis, lipid accumulation and disturbed regeneration [14] which are all hallmarks of the DMD phenotype further supporting the cyclical nature and mutual dependence.

Gerber’s revolutionary work on the circadian regulation of the RhoA-actin-SRF pathway, eloquently describes how the hypothalamic suprachiasmatic nucleus (SCN), transmits systemic cues thereby activating RhoA in peripheral tissues [8]. They go on to demonstrate that diurnal polymerisation (F-actin) and de-polymerisation (globular (G)-actin) of actin influences SRF expression and transcription of specific downstream circadian targets (*Per1* and *Per2*) and output genes (*Nr1d1* and *Rora1*) [8, 9]. However, disruption of this pathway by removing alternative upstream proteins intrinsic to the cascade has not been shown. Given the proximity of dystrophin to F-actin (which it specifically binds and stabilises) combined with its integral relationship with SRF transcription, we hypothesise that dystrophin loss leads to a shift in actin de-polymerisation, which affects SRF expression and downstream circadian gene expression, thus resulting in circadian dysregulation in DMD models.

## Results

To assess whether dystrophin loss disrupts the RhoA-actin-SRF pathway, an *in vitro* assay was designed using siRNAs targeting the *Dmd* gene to down-regulate dystrophin protein in a skeletal muscle cell line (H2K 2B4 myoblasts) [15]. Differentiated myotubes were transfected twice with 100nM siRNA, and collected 49 hours following the second transfection, thereby representing circadian time 1 (CT1). Efficient transfection resulted in undetectable amounts of dystrophin protein (Fig 1A) and markedly reduced *Dmd* transcript levels (Fig 1B). Dystrophic conditions resulted in altered RhoA activation (Fig 1C) and decreased F/G-actin ratios (Fig 1D). To illustrate circadian oscillation patterns of core clock genes involved in the transcriptional auto-regulatory feedback loop following abrogation of dystrophin, cells were collected every 4 hrs over a 24 hr period, and indicate significant alterations (Fig 1F). *Per1*, *Per2* and *Clock* expression were significantly down-regulated at certain time points, whilst *Cry2* and *Arntl1* were upregulated. Expression of other downstream SRF target genes in the actin-cascade, *Nr1d1* and *Acta [9],* were also significantly lower in dystrophin deficient samples, and interestingly *Srf* expression was significantly upregulated. Whilst *Srf* upregulation was unexpected, this may be due to activation of alternative compensatory pathways such as the ternary complex factor (TCF) family of Ets domain proteins (MAPK pathway) which regulates transcription of growth responsive genes [9]. Importantly, *Srf* upregulation does not denote SRF activation via the RhoA-actin pathway, which would require nuclear translocation of MRTF. SRF chromatin-immunoprecipitation (ChIP) combined with qRT-PCR revealed reduction of occupancy of SRF on the *Nr1d1* regulatory region and partial reduction on *Per2*, which are two established targets of the MRTF-SRF cascade [9], thereby indicating impaired SRF signalling (Fig 1G, see Fig S1 for SRF-binding motifs). Note: other clock genes do not contain SRF-binding motifs so were not investigated by ChIP [9]. In order to investigate to what extent dystrophin loss in human DMD patients resulted in similar biochemical events, we obtained DMD muscle biopsies from patients (tissues collected between 8-10am in the morning and patients fasted from midnight the evening before biopsy). Interestingly, a wide array of mutations showed down-regulation of key RhoA-actin-SRF targets *PER1*, *PER2* and *NR1D1* in nearly all samples (Fig 1H).

**Figure 1.**
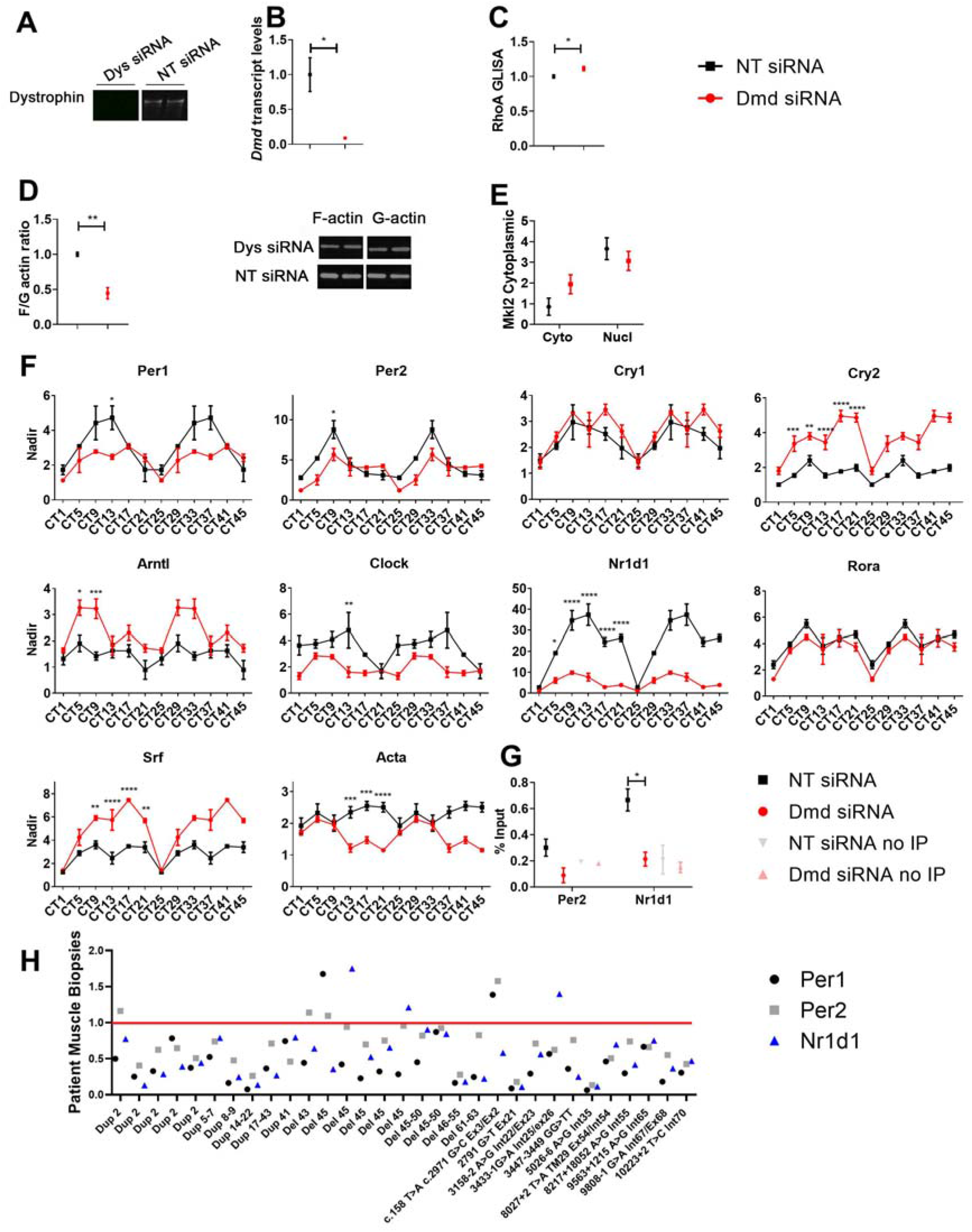
Disruption of RhoA-actin-SRF pathway in dystrophic cells and patient muscle biopsies. To mimic dystrophic conditions, H2K 2B4 myotubes were transfected with siRNAs targeting the *Dmd* gene (Dmd siRNA, red), and a non-targeting (NT siRNA, black) siRNA was used for control. **(A)** Dystrophin was successfully down-regulated at protein, and **(B)** transcript level. Additionally, the absence of dystrophin resulted in **(C)** altered RhoA activity, **(D)** reduced F/G-actin levels, and **(E)** lower nuclear MRTF accumulation (TBP and β-actin for loading controls). For gene transcript studies, cells were placed is serum free media and collected over a 24 hour time course (double plotted-48 hours- to show oscillation pattern). **(F)** Circadian time (CT) gene expression data indicates diurnal oscillation patterns and down-regulation of pertinent core clock genes (*Per1* and *Per2*) and down-stream targets for RhoA-actin-SRF (*Nr1d1* and *Acta1*; housekeeping gene-*Gapdh*). **(G)** SRF chromatin immunoprecipitation (ChIP) revealed reduced enrichment of down-stream targets, *Per2* and *Nr1d1*. **(H)** Muscle biopsies were obtained from an array of patients with different mutations or deletions in the dystrophin genes, and indicate down-regulation of RhoA-actin-SRF target genes in most cases (housekeeping genes-*Rpl13a*). All samples were normalised to a pooled skeletal muscle sample from healthy volunteers, as indicated by red line. For RhoA GLISA, F/G actin ratio and MRTF westerns, data was normalised to NT control (n=3; Two-tailed Student’s *t*-test). For CHIP n=3; two-tailed Student’s *t*-test. For RT-qPCR CT data, the nadir was determined as the minimum value across both treatments (Dmd and NT siRNAs) and CTs and applied to all samples; nadir normalised to 1 (n=2-3 for each siRNA and time point; Two-way ANOVA with Bonferroni *post-hoc* test performed). Mean values reported with SEM; ***p⍰<⍰0.001, **p⍰<⍰0.01, *p⍰<⍰0.05.

Given these *in vitro* and biopsy results, we predicted that SRF signalling in peripheral muscle would also be interrupted in dystrophic mice, leading to molecular deficits. The dystrophin-utrophin knockout (dKO) model presents with a severe phenotype that closely recapitulates disease in patients, specifically severe progressive muscular dystrophy, premature death and a plenitude of physiological and molecular aberrations [16]. The oscillation patterns of core clock genes in the *tibialis anterior* (TA) muscle of 5 week old male mice were assessed over a 24 hour period (double plotted). This model revealed profound alterations in core clock gene expression, with markedly similar patterns to that observed in the dystrophic-H2K 2B4 myotubes. The amplitude of SRF-target genes *Per1*, *Per2* and *Nr1d1* was lower in 5 week old dKO animals compared to C57BL10 (Fig 2A), suggesting dampened rhythmicity. Gene expression of *Cry1, Cry2, Arntl, Clock* and other downstream targets (*Acta* and *Rora1*) were all significantly upregulated at various ZTs. As observed in the H2K 2B4 model, *Srf* expression was again upregulated under dystrophic conditions. Additionally, F/G-actin ratios (Fig 2B), and MRTF (Mkl2) nuclear-fraction protein levels (Fig 2C) were lower in 5 week old dKO animals compared to control animals. Interestingly, MRTF cytoplasmic-fraction protein levels were also lower in dKO animals, which suggests less MRTF is present in dKO animals. It is important to note that both F/G actin ratios and MRTF levels exhibited significant diurnal changes in skeletal muscle. These results, in combination with the *in vitro* data, confirm that the actin-pathway is indeed perturbed in muscle due to dystrophin loss, and displays diurnal alterations.

**Figure 2.**
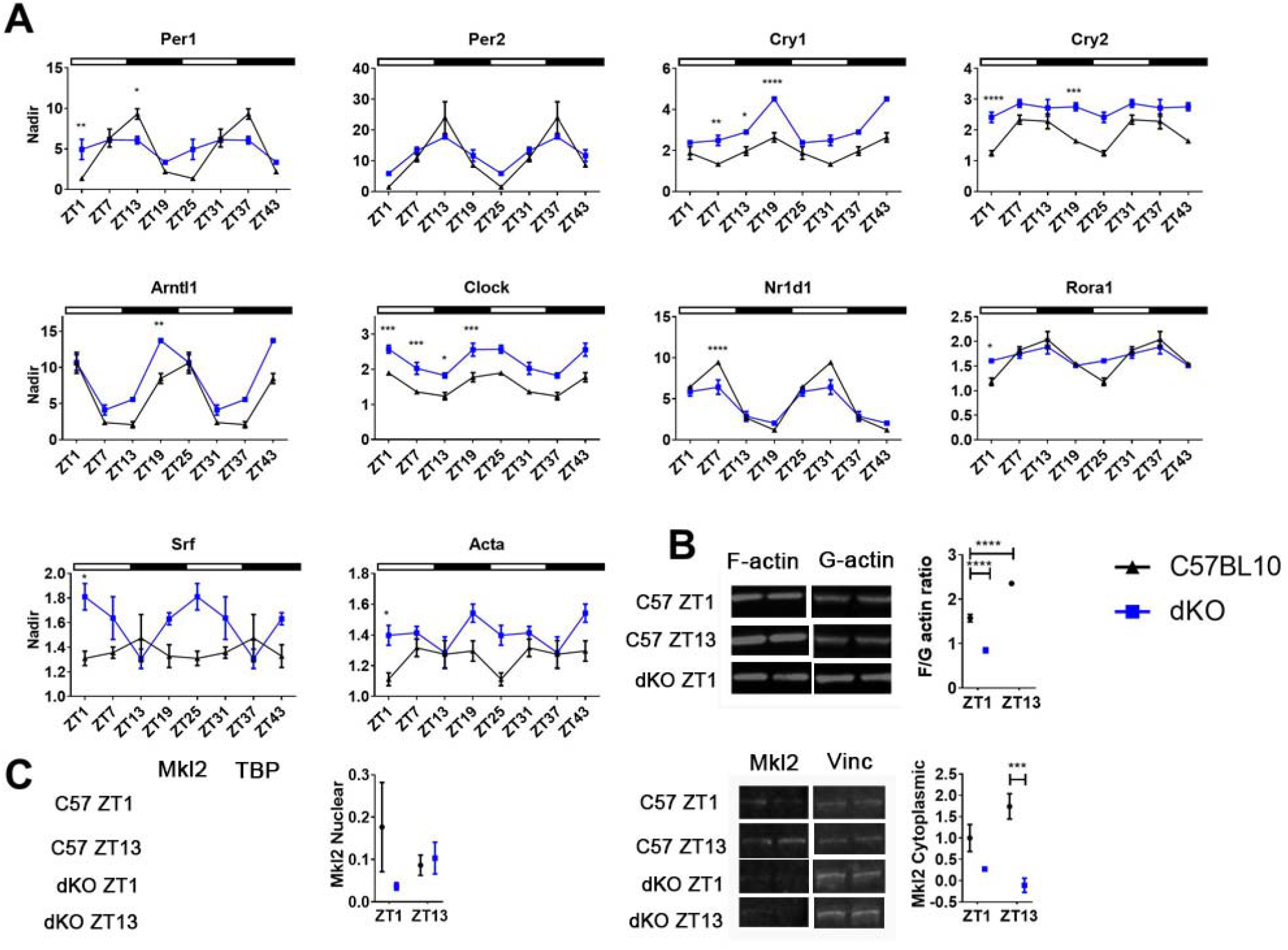
Diurnal changes in RhoA-MRTF-SRF cascade components in *tibilias anterior* of dystrophin-utrophin (dKO) mouse model. Tissues were collected over a 24 hour time course and double plotted (48 hours) to better **(A)** illustrate the oscillation pattern of core clock genes in *tibilias anterior* (TA) of 5 week old *dKO* animals which were significantly altered compared to C57BL10 (Zeitgeber-ZT). Down-stream targets for RhoA-actin-SRF pathway (*Nr1d1*, *Rora1* and *Acta*) and indeed *Srf* were also altered (*Gapdh* used as housekeeping gene). **(B)** F/G-actin protein ratios in *dKO* mice are significantly down-regulated, **(C)** as were nuclear and cytoplasmic MRTF fraction levels (TBP and vinculin used as loading controls). In addition C57BL10 animals exhibit diurnal changes in F/G actin ratio and MRTF fraction levels. Light and dark periods represented by outlined (light) and solid bars (dark). For RT-qPCR ZT data, the nadir was determined as the minimum value across both genotypes and ZTs and applied to all samples; nadir normalised to 1 (n=3-4 for each for each genotype and time-point). For F/G actin ratio and MRTF westerns, data was normalised to C57BL10 (n=3-4). One or Two-way ANOVA with Bonferroni *post-hoc* test performed. Mean values reported with SEM; ***p⍰<⍰0.001, **p⍰<⍰0.01, *p⍰<⍰0.05.

We further anticipated that systemic cues from the central clock (SCN) to SRF in peripheral muscle would be interrupted. Whilst the dKO mouse model closely recapitulates the dystophic phenotype in patients and is a good molecular model for DMD, it’s severe phenotype including reduced lifespan (approx. 5-8 weeks) and marked reduction in activity, preclude extensive locomotor behaviour studies. As such, the less affected dystrophic model, *mdx*, was utilised for the extensive battery of locomotor tests. We show for the first time that dystrophin protein is expressed in the SCN of C57BL10 mice but not *mdx* animals (Fig 3A), and therefore it was pertinent to assess whether there were any obvious abnormalities in the circadian locomotor behaviour in dystrophic mice. Wheel-running activity of 20 week old (symptomatic) male *mdx* and C57BL10 mice were recorded under various conditions, and representative actograms under 12hr:12hr light-dark (LD), 24hr dark (DD), and 24hr light (LL; Fig 3B) are shown. Under normal LD conditions, no significant differences in the number of bouts or total activity between mouse cohorts was observed, indicating that the endurance level of *mdx* mice was normal (Fig 3C). This observation was valuable for the interpretation of subsequent data, as it indicated differences between genotypes was not due to degenerative repercussions of the dystrophic phenotype but rather signalling cues from the SCN. Interestingly, during the light phase of LD, activity of *mdx* mice was markedly reduced, and they exhibited delayed onset into dark phase (phase angle). Following 6-hour phase advance bouts, *mdx* mice were capable of re-entraining to the shifted cycle in a similar manner to control animals (Fig 3D). Animals were then placed in DD, where *mdx* animals again indicated a delayed onset (phase angle) on release into dark (Fig 3E), suggesting that their endogenous clock may be out of phase. Again endurance of *mdx* animals was maintained during DD and they ran for a similar period compared to C57BL10 animals, however their free running period was significantly shorter (Fig 3F; *mdx* run half an hour shorter). During the DD phase, mice received a light pulse 4-hours after they started exercise (CT16), and *mdx* mice displayed no difference in the ability to shift the clocks phase in response to this nocturnal light compared to C57BL10 animals (Fig 3G). In light of our altered dystrophin-associated-RhoA-actin-SRF cascade hypothesis, altered activity may be due to dystrophin loss in the SCN leading to alterations in core clock gene, *Per2,* which has been associated with shorter circadian period and loss of circadian rhythmicity in constant darkness [17]. The delayed phase angle in LD and DD, combined with extreme lack of activity during LL suggest a severe aversion to light stimuli in *mdx* animals. Long-term exposure to light has been shown to affect neurons in the SCN and reduce rhythmicity [18], which aggravated by the loss of dystrophin in the SCN, may explain the considerable changes in activity of *mdx* mice. Altogether, these data demonstrate the SCN is profoundly affected in dystrophic mice.

**Figure 3.**
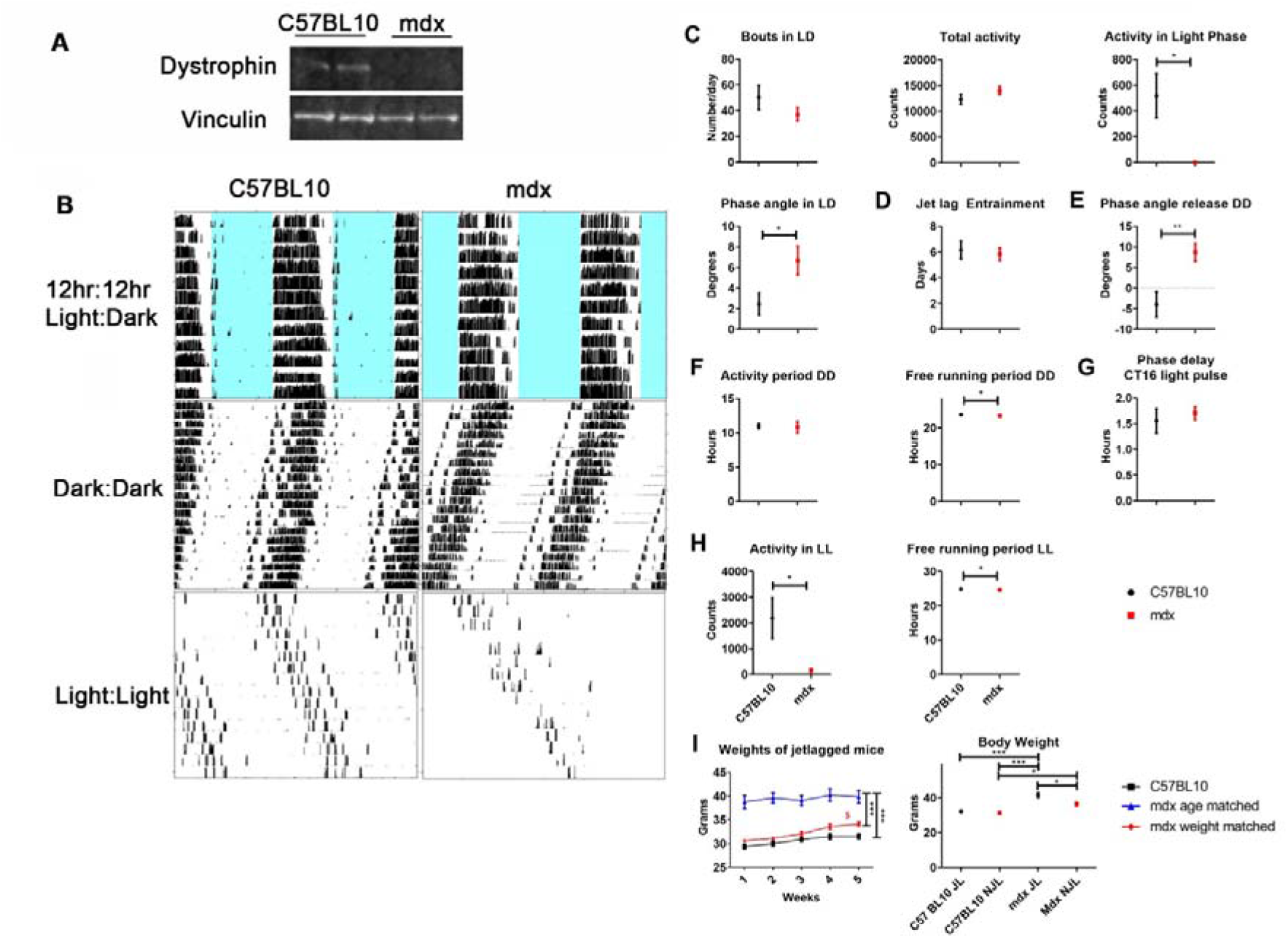
Perturbed circadian rest-activity and response to jet-lag in dystrophic mouse model. **(A)** Dystrophin western blot shows the absence of protein in the SCN of *mdx* animals (vinculin used as loading control). Locomotor behaviour of *mdx* and C57BL10 mice were assessed and representative wheel-running activity blots **(B)** shown; 12h:12h light:dark cycle (LD), dark:dark (DD) and light:light (LL). Data analysed and represented as graphs: **(C)** behaviour during LD, **(D)** following 6-hour phase advancements, **(E)** phase angle on release into DD, **(F)** behaviour during DD, **(G)** phase delay following a light pulse 4-hours post exercise commencement, **(H)** and behaviour during LL. **(I)** Weights of mice undergoing 5 week jetlag protocol assessed (C57BL10, *mdx* age matched and *mdx* weight matched), as well as total body weight of jetlagged (JL) and non-jetlagged (NJL) *mdx* and C57BL10 cohorts. Student’s *t*-test for comparisons between 2 groups (Two-tailed). For weights over 5 weeks of jetlag: black asterisk on right-one-way ANOVA comparing all 3 cohorts; dollar sign-one way ANOVA with Bonferroni *post-hoc* between each cohort at each time point. For body weights with JL and NJL, One-way ANOVA with Bonferroni *post-hoc* test performed (For wheel running activity *n*=5-6, jetlag study *n*=3-4; ***p⍰<⍰0.001, **p⍰<⍰0.01, *p⍰<⍰0.05).

Male *mdx* and C57BL10 mice were also subject to repeated bouts of chronic jetlag (JL) whereby lighting conditions were advanced 6 hours every week for 5 weeks (mice weighed weekly). Age-matched *mdx* were significantly heavier than C57BL10 mice (15 weeks old at start of protocol, 22 weeks by end), therefore a ‘weight matched’ group (12 weeks at start, 20 weeks at end) was also assessed. ‘Weight matched’ *mdx* mice significantly increased in weight over 5 weeks, whereas there was no increase in weight in the C57BL10 cohort (Fig 3I). The JL cohorts were further compared to non-jet lagged (NJL) groups. No significant differences were observed between JL and NJL C57BL10 cohorts (all mice 22 weeks of age). Although NJL *mdx* animals were significantly heavier than both C57BL10 cohorts, most importantly the JL *mdx* cohort was significantly heavier than all other groups. Jetlag causes changes in phase of entrainment in the SCN and peripheral clock [19]. Some clocks entrain faster than others which can cause internal desynchrony. If the synchronising signal such as SRF is lost (in this case due to disruption of RhoA-actin-SRF cascade), it is anticipated there will be a greater disruption and desynchrony from jetlag protocols leading to changes in weight. This would account for why the C57BL10 animals resisted weight gain under the short JL conditioning protocol, whereas the *mdx* animals do. Together these data indicate altered signalling between the SCN and peripheral tissues in dystrophin-deficient *mdx* mice.

As locomotor experiments were performed in the *mdx* model, gene expression patterns of core clock genes in the TA muscle of 20 week old male mice were assessed over a 24 hour period (double plotted). A shift in phase was observed for *Per1*, *Per2*, *Cry1, Cry2* and *Arntl1,* resulting in significant differences in gene expression at certain time-points within the day (Fig 4). For instance, the expression of *Per2* in *mdx* mice peaked at ZT5, but in C57L10 animals peaked at ZT9. Gene expression of downstream SRF target, *Rora1,* was significantly down-regulated (ZT1, ZT17 and ZT21), and minor alterations in *Nr1d1* (ZT21) and *Acta1* (ZT1) were observed.

**Figure 4.**
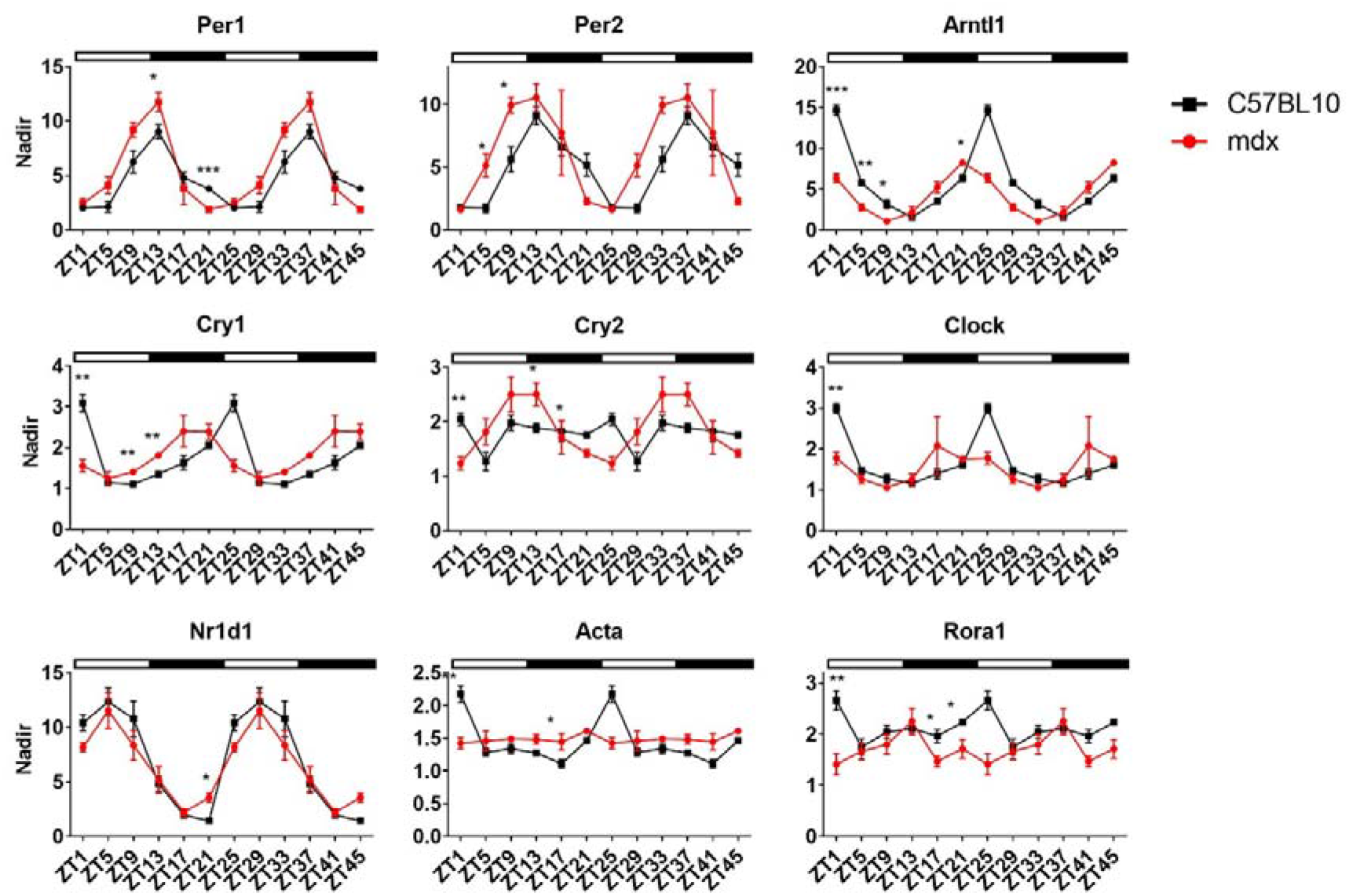
Altered expression of clock genes in *tibilias anterior* of *mdx* mouse model. Tissues were collected over a 24 hour time course and double plotted (48 hours) to better illustrate the oscillation pattern of core clock genes in *tibilias anterior* (TA) of 20 week old *mdx* animals which were significantly altered compared to C57BL10 (Zeitgeber-ZT). Down-stream targets for RhoA-actin-SRF pathway (*Nr1d1*, *Acta* and *Rora1*) were also altered (*Atp5b* used as housekeeping gene). Light and dark periods represented by outlined (light) and solid bars (dark). The nadir was determined as the minimum value across both genotypes and ZTs and applied to all samples; nadir normalised to 1 (n=3-4 for each for each genotype and time-point; Two-way ANOVA with Bonferroni *post-hoc* test performed). Mean values reported with SEM; ***p⍰<⍰0.001, **p⍰<⍰0.01, *p⍰<⍰0.05.

Most importantly we observe alterations in integral components of the RhoA-actin-SRF cascade (in particular SRF targets and F/G-actin ratios) in all dystrophic models described. Whilst the *mdx* gene dataset differs to the dKO and H2K 2B4 myotube models, we regard the dKO model with greater esteem given its phenotype and correlation with patient disease progression [16], which is supported by the biopsy data (Fig 1H). It further correlates closely with cell-culture data in which dystrophin is specifically abrogated (Fig 1F). However, in order to illustrate locomotive aberrations, and systemic cues with the SCN, it was imperative we look in the milder *mdx* model. The remarkable lifespan and generally mild phenotype of *mdx* mice is poorly understood, but it is likely due to multiple compensatory events that ensue and may account for the variances observed between the dystrophic models. As multiple inputs can regulate the clock, it is difficult to predict how the clock will react when one input is removed. *In vivo* complexities in the form of protein or signalling interactions with serum, hormones (particularly glucocorticoids as *Per2* has a glucocorticoid receptor-binding site) and neurological signals or mechanisms triggered to compensate for disturbed RhoA-actin-SRF pathway may be involved. One such compensatory mechanism is upregulation of utrophin, a homologue of dystrophin, which is knocked out in dKO but present in *mdx* mice. Utrophin has been shown to bind and maintain F-actin polymerisation and therefore seems an obvious candidate in the RhoA-actin-SRF signalling cascade [20]. We show the presence of utrophin gene and protein expression in *mdx* mice (Fig S2A), however in H2K 2B4 myotubes, *Utrn* gene expression and protein levels were reduced (Fig S2B). Similarly, in DMD patient biopsies, *UTRN* gene expression was down-regulated in all samples with the exception of one, which incidentally also showed upregulation of *PER1* and *PER2* for that sample (Fig S2C). Indeed, linear regression analysis of *UTRN* expression versus *PER1* and *PER2* indicate a significant correlation pattern (*PER1* R= 0.1687, *PER2* R= 0.4542; Fig S2D). Thus, utrophin may compensate for dystrophin loss in *mdx* animals, and loss of both utrophin and dystrophin may lead to greater F-actin instability and downstream SRF activity, as observed in the H2K 2B4 and dKO data.

To assess the RhoA-actin SRF cascade further, and determine whether abrogation of other upstream components of the pathway, such as actin, or indeed SRF itself, results in similar changes in the expression of target genes, siRNAs were used to specifically knock-down *Srf* and *Acta* in H2K 2B4 myoblasts. In order to compare with dystrophin knock-down experiments, differentiated myotubes were transfected twice with 100nM *Srf* and *Acta* siRNAs, alongside *Dmd* transfected myotubes, and collected 49 hours following the second transfection, thereby representing CT1. Myotubes were also treated with lower concentrations of siRNA to confirm gene expression was stable and that myotubes were healthy (Fig S3). Interestingly, *Srf*, *Acta* and *Dmd* down-regulation, appear to reciprocally modulate each other resulting in lower expression of all genes for all cohorts ie. *Srf* knock-down results in lower *Dmd* and *Acta* gene expression and *vice versa* (Fig 5). Additionally, all siRNA treatment groups resulted in reductions of RhoA-actin-SRF target genes, *Per1* and *Per2*, and *Nr1d1* and *Rora*. Together, this illustrates how intertwined and mutually dependent these genes are in maintaining homeostatic balance of the RhoA-actin-SRF cascade.

**Figure 5.**
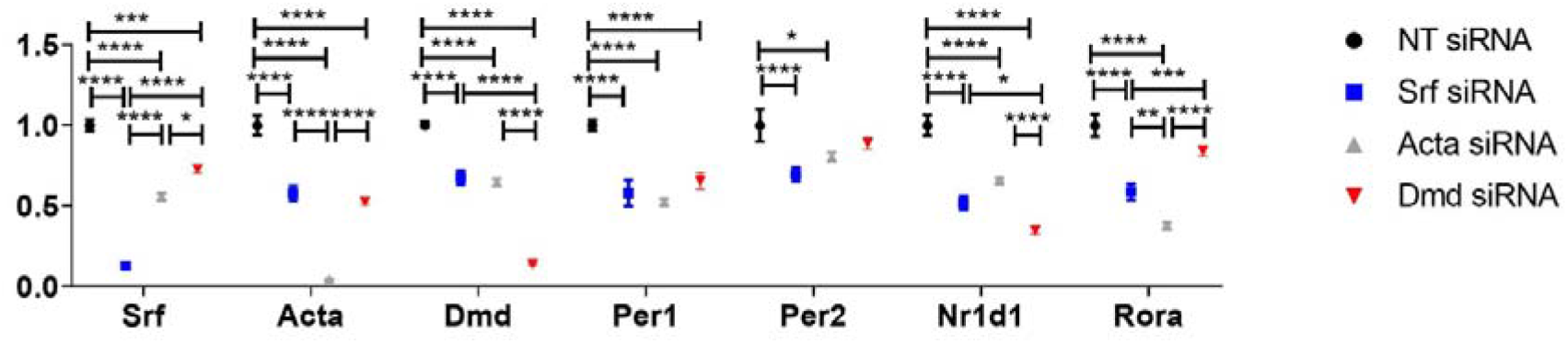
Abrogation of alternative RhoA-actin-SRF components leads to a reduction in the expression of SRF target genes. H2K 2B4 myotubes were transfected with 100nM siRNAs targeting the *Srf, Acta* and *Dmd* genes (*Srf* siRNA, blue, *Acta* siRNA grey and *Dmd* siRNA, red), and a non-targeting (NT siRNA, black) siRNA was used for control. Gene expression data indicates successful knock-down of respective genes and down-regulation of pertinent core clock genes (*Per1* and *Per2*) and other down-stream targets for RhoA-actin-SRF (*Nr1d1* and *Acta1*; housekeeping gene-*Gapdh*). Data normalised to NT siRNA; n=3; Two-way ANOVA with Bonferroni *post-hoc* test performed. Mean values reported with SEM; ***p⍰<⍰0.001, **p⍰<⍰0.01, *p⍰<⍰0.05.

## Discussion

Due to the severe repercussions of genetic disorders, many events are often overlooked, as efforts are primarily aimed at targeting the underlying genetic defect. This is most certainly the case for Duchenne muscular dystrophy (DMD), a monogenic disorder resulting in muscle wasting and cardiomyopathy in affected boys. Indeed these boys experience abnormalities in sleeping patterns [21], and nocturnal hypoxaemia and hypercapnia [22] which may be attributed to the dystrophic phenotype-specifically the deterioration in respiratory muscles. Here we propose this may also be due to a circadian deficit.

We show circadian perturbations in a number of dystrophic models and suggest a mechanistic rationale for these changes by investigating components of the RhoA-actin-SRF cascade. This seemed a logical approach given F-actins interaction with dystrophin, and its importance in the RhoA-actin-SRF cascade, which is crucial for muscle homeostasis. In the case of healthy muscle, RhoA regulates polymerisation (F-actin) and de-polymerisation (G-actin; Fig 6). During de-polymerisation G-actin preferentially binds to MRTF, but when this shifts to the polymerisation phase the G-actin pool diminishes and unbound MRTF translocates to the nucleus and influences SRF expression. SRF activation in turn regulates transcription of target genes-*Per1, Per2*, *Rora1*, *Nr1d1*, and *Acta*. Here we propose that the absence of dystrophin reduces F-actin levels resulting in shifts to the de-polymerised state. G-actin binds to MRTF, which we show remains cytosolic thereby hampering SRF activation (Fig 6). Additionally, we show for the first time that dystrophin is absent in the SCNs of dystrophic mice, and that these animals exhibit behavioural alterations, indicative of disrupted central circadian signalling within the SCN. As such, the lack of dystrophin results in circadian disruption that manifests with physiological and molecular alterations in dystrophic models.

**Figure 6.**
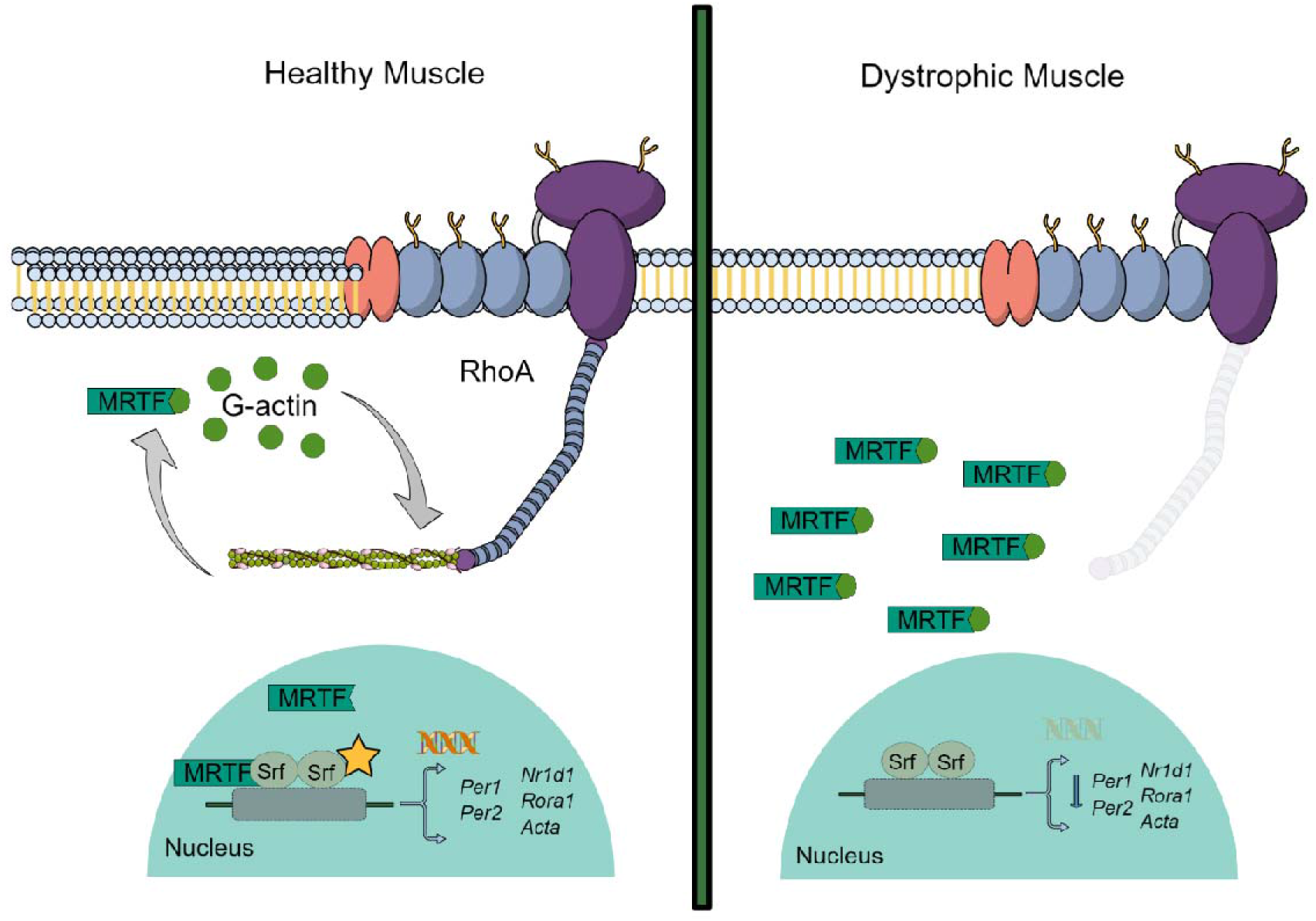
Schematic illustrating RhoA-actin-SRF signalling pathway. In healthy muscle, systemic cues activate RhoA in peripheral tissues which in turn regulates diurnal polymerisation (F-actin) and de-polymerisation (globular (G)-actin) of actin. During the de-polymerisation stage, G-actin preferentially binds to myocardin-related transcription factor (MRTF). However, when this shifts to the polymerisation phase, the G-actin pool diminishes and unbound MRTF translocates into the nucleus and influences expression of serum response factor (SRF). SRF activation regulates transcription of SRF target genes including the core clock genes-*Per1* and *Per2*, as well as secondary loop clock genes-*Rora1* and *Nr1d1*, and cytoskeletal genes such as *Acta*. When dystrophin is absent, F-actin is not stable and the cell shifts to the de-polymerised state. G-actin binds to MRTF and therefore SRF transcription is hampered.

Given that dystrophin regulates circadian signalling in peripheral tissues, this suggests that related dystrophin glycoprotein complex (DGC) proteins may be implicated in dystrophin and F-actin tethering, loss of which could also result in circadian dysregulation. Indeed this is supported by knock-down of *Srf* and *Acta,* alternative components of the RhoA-actin-SRF pathway, which also lead to down-regulation of target genes. Thus, muscular dystrophy disorders, such as limb-girdle muscular dystrophy[23], in which DGC and sarcomeric proteins are affected should be assessed for circadian abnormalities.

It is possible that some pharmacological interventions used to improve the muscle phenotype in DMD patients, may have inadvertently modulated circadian rhythm resulting in physiological improvements, such as observed with the use of melatonin [24] and glucocorticoids [25]. These hormones oscillate throughout the day, are highly regulated by sleep-wake cycles, and are governed by SCN signalling [26, 27]. This is particularly interesting given that dystrophin is absent in the SCN and may have consequential repercussions on endocrinological processes in DMD patients. It has also been shown that the glucocorticoid, dexamethasone, induces the transcription factor, *Klf15*, with beneficial ergonomic effects on dystrophic muscle [28]. Klf15 regulates a multitude of processes including metabolism [29, 30] and nitrogen homeostasis [31], but most importantly does so in a circadian fashion, and has also been shown to be altered in another neuromuscular disorder, spinal muscular atrophy [32]. Dystrophic mice were adversely affected by constant light exposure. LL causes arrhythmicity in the long-term, and short-term exposure is a method used for period lengthening which is a common feature of the clock in nocturnal rodents. It is uncertain what effect LL would have in human subjects, but constant dim light has been used (rather than DD) to unmask the central clock, suggesting light augmentation may be an appropriate therapy for DMD patients. Other therapies including exercise, dietary modification or drugs to mitigate disease pathology (such as lower calcium) [33] may also usefully be investigated in the context of DMD treatment. This study therefore reveals both a hitherto an unanticipated role for dystrophin in peripheral muscle tissues and novel avenues for further research and possible therapeutic intervention applicable to a range of muscular dystrophy disorders.

## Materials and Methods

The datasets generated during this study are available from the corresponding author on reasonable request.

### *In vitro Dmd* knock-down

H2K 2B4 myoblasts were cultured in flasks coated with Matrigel (Corning) and in DMEM culture media supplemented with 2% chick embryo extract (Seralabs), 10% fetal calf serum (Gibco) and 1% antibiotics (Gibco); 33⁰C and 10% CO_2_. Myoblasts require differentiation into myotubes in order to produce dystrophin, and therefore differentiation medium containing 5% horse serum (Gibco) was used (DMEM supplemented with 1% antibiotics). Dystrophin was knocked-down using siRNAs targeting the dystrophin transcript (Dharmacon; see Table S1). H2K 2B4 cells were transfected with siRNAs (100nM) and lipofectamine RNAiMAX (ThermoFisher Scientific) at day 0 and day 2 of the differentiation process. For RhoA GLISA, F/G actin and MRTF studies, cells were harvested 49hrs later, on day 4. Experiments were performed in duplicate, whereby cells were scraped, pooled and further split into 3 vials for downstream analyses (ie. protein, F/G-actin). This experiment was repeated 3 times to attain 3 biological replicates. Data shown relative to control siRNA for each experiment. For RT-qPCR study, 49hrs after second transfection, media was removed, cells were washed with PBS, and serum free media (DMEM supplemented with 10mM HEPES and 2% B27-Gibco) was placed in wells. Cells were collected an hour later (CT1), and then every 4hrs over a 24hr period. Note: cells were not synchronised with dexamethasone due to the known pharmacological effects observed in dystrophic cells which would likely result in complicated interpretation of the results. For the ChIP experiment, cells were grown on 15cm plates. Plates were transfected with 100nM siRNA using lipofectamine RNAiMAX at day 0 and day 2 of the differentiation process. Two days later cells were cross-linked with 36.5% formaldehyde and quenched with 1M glycine. Cells were collected by scraping in cold PBS, and following centrifugation, cells were resuspended in CLB and incubated for 10mins. Nuclei were pelleted and frozen at −80 degrees.

### ChIP

ChIP was performed using Dynabeads Protein G (ThermoFisher, 1003D) and 5μg of anti-SRF antibody (ab53147; Abcam) as previously described [34, 35]. The immunoprecipitated (SRF antibody in immunoprecipitation (IP) reaction), control (beads only) and input DNAs were used as templates for qPCR, with SYBR Green PCR Master Mix (Applied Biosystems) and gene-specific primers (Table S1). The % Input was calculated for each gene region by subtracting the input from the IP sample (delta Ct) and then performing a power calculation (100×2^ (ΔCt)). The negative background signal (as determined by the ‘no antibody’ control) was calculated in a similar manner.

### SRF and actin knock-down

H2K 2B4 myoblasts were again cultured in matrigel coated flasks using DMEM culture media (2% chick embryo extract, 10% fetal calf serum and 1% antibiotics; 33⁰C and 10% CO_2_). Myoblasts were differentiated using medium containing 5% horse serum (Gibco) in DMEM supplemented with 1% antibiotics. Myoblasts were transfected with *Srf* and *Acta* siRNAs (Dharmacon; see Table S1) using lipofectamine RNAiMAX (ThermoFisher Scientific) at day 0 and day 2 of the differentiation process. Cells were harvested 49hrs after the second transfection, on day 4.

### RNA extraction and RT-qPCR

For patient biopsies, informed consent was obtained from DMD patients’ for standard diagnostic purposes, including muscle biopsy. In addition, we obtained informed consent to use muscle biopsies for research activities, according to the Telethon Project N. GGP07011, Ethical approval N. 9/2005, 25 October 2005, by the S.Anna University Hospital Local Ethical Committee, Italy. Patients fasted overnight before the biopsy, which was performed at 8-10am. The control RNA was obtained commercially and consists of multiple human skeletal muscle samples pooled together (Ambion). RNA was extracted using an RNAeasy kit from Qiagen as per manufacturer’s instructions.

For mouse muscle and myotubes, total RNA was extracted from tissue or cells using Trizol (Thermo Scientific Technologies).

All RNA was reverse transcribed using a High Capacity cDNA synthesis kit (Applied Biosystems). The cDNA was run using TaqMan probes sets (Applied Biosystems) and gene specific primer sets for the core clock genes (Integrated DNA Technologies; see Table S1) on the StepOne Real-Time PCR system (Applied Biosystems). Housekeeping genes were determined by running samples from patients (Human geNorm kit; Primer Design), cell culture and each mouse model (Mouse geNorm kit; Primer Design) with primers from geNorm qRT-PCR normalisation kit for. *Rpl13a* was the most stable house-keeping gene for Patient biopsies, *Gapdh* for cell culture and dKO samples, and *Atp5b* was the best for *mdx* samples. Tissues were collected over a 24 hour time course, and data was double plotted (48 hours) to better illustrate the oscillation pattern of core clock genes in muscle. The nadir was determined as the minimum value across all genotypes and ZT’s and applied to all samples; nadir normalised to 1.

### RhoA-activity quantification

RhoA activity was calculated using a RhoA G-LISA Activation Assay kit as per manufacturer’s instructions (Cytoskeleton). Briefly, cells were washed, lysed and snap-frozen until required. Protein quantification was performed using Precision Red and samples were run on Rho-GTP ELISA.

### F/G actin westerns

Actin protein levels were calculated using a G-actin/F-actin *in vivo* Assay Biochem Kit as per manufacturer’s instructions (Cytoskeleton). Briefly, samples were lysed and underwent ultra-centrifugation to separate fractions. The G-actin supernatant was removed and the F-actin was resuspended in F-actin depolymerisation solution. Samples were run on 10% Bis-Tris gels (Thermo Scientific Technologies) and blotted as per supplier’s instruction.

### MRTF westerns

Nuclear and cytoplasmic fractions were collected from tissue culture samples and dKO tissues using the NE-PER™ kit as described in manufacturer’s instructions (ThermoFisher Scientific). Samples were run on 10% Bis Tris gels (Thermo Scientific Technologies), blotted onto PVDF membrane (Millipore) and probed with Mkl2 (ab191496; Abcam). For nuclear fractions TBP (ab51841; Abcam) and U2AF65 (ab37530; Abcam) were used as loading controls, and for cytoplasmic fractions vinculin (hVIN-1; Sigma) and β-actin were used as loading controls. Blots were visualised using IRDye 800CW goat-anti rabbit IgG or IRDye 800CW goat-anti mouse IgG (Licor) on the Odyssey imaging system.

### Dystrophin/Utrophin protein quantification

Dystrophin and utrophin protein levels were quantified as previously described [36]. Briefly, homogenised samples were run on 3-8% Tris-Acetate gels (Thermo Scientific Technologies), blotted onto PVDF membrane (Millipore) and probed with DYS1 (Novocastra, dystrophin) or MANCHO antibody (KED laboratory-Oxford, utrophin). Blots were visualised using IRDye 800CW goat-anti mouse IgG (Licor) on the Odyssey imaging system. For dystrophin, vinculin (hVIN-1; Sigma) was used as loading control, and for utrophin samples were quantified against total protein using Coomassie stain (Sigma).

### Animals housing

All procedures were authorized and approved by the University of Oxford ethics committee and UK Home Office (project licence PDFEDC6F0, protocols 19b1, 2 and 4). Investigators complied with relevant ethics pertaining to these regulations. Procedures were performed in the Biomedical Sciences Unit, University of Oxford. Mice were allowed food and water *ad libitum*.

### *In vivo* experiments for gene expression, jetlag and wheel-running studies

All experiments were performed in male dystrophic C57BL/10ScSn-*Dmd^mdx^*/J (mdx; JaxLabs), C57B/L10ScSn-Utrn^tm1Ked^ Dmd*^mdx^*/J (dKO; JaxLabs) and control C57BL/10ScSnJ (C57BL10; Envigo and Oxford BSB) mice. For the 24 hour ZT study, mice (5 week old dKO, littermate *mdx* and C57BL10, or 20 week old *mdx* and C57BL10) were housed under a strict 12:12hr light-dark cycle (LD; 400 lux white light) for 2-3 weeks, after which mice were sacrificed every 4-6 hrs over a 24 hr time course (n=3-5 for each genotype and timepoint). Briefly, mice were culled by cervical dislocation and eyes removed. Animals during the night cycle were culled under dim red light. The TA was dissected, snap frozen and stored at −80°C.

For wheel-running experiments, each mouse was placed in a large cage fitted with a running wheel (n= 6 for each genotype). Activity was recorded using the *Clock*Lab console. Mice were first entrained to a 12:12hr LD cycle at 400lux white light for 2 weeks, and then underwent 6 hour phase advance entraining, before being placed into constant darkness (DD).

For the jetlag weight study, animals were group housed under a normal 12:12 LD cycle (400 lux white light) for two weeks to establish stable entrainment (n= 3-4 for each genotype and condition). The LD cycle was then advanced by 6h every week for 5 weeks. Animals were weighed close to ZT13 each week. Animals were culled at the end of the protocol and tissues harvested, weighed and snap frozen.

### Statistics

All statistics were performed using SPSS or GraphPad. For experiments including 3 or more comparisons, One-way or Two-way ANOVA was performed with either Bonferroni or Games-Howell *post-hoc* corrections. For the weight gain experiments, a repeated measures One-way ANOVA was used to investigate the overall weight gain differences between the groups. For experiments between only 2 groups, student’s *t*-test (Two-tailed) was performed.

## Acknowledgements

We would like to thank the BMS facility at Oxford for their care and support of the animals, and Dr Peter Oliver for the use of the controlled light housing units. We would also like to thank Dr Ben Edwards and Prof. Kay Davies for providing utrophin antibody. We thank the Telethon Italy Network of Genetic Biobanks (Dr. Marina Mora) for providing biological samples. Special thanks are due to Duchenne Parent Project Italia Onlus for granting AF and MSF DMD diagnostic studies, and Duchenne Parent Project Italy General Grant for DMD cell biobank funding. CB is supported by the British Heart Foundation and Muscular Dystrophy UK. TvW was supported by Muscular Dystrophy UK. MB was an SMA Trust Career Development Fellow when at the University of Oxford. KEM was funded by the MDUK and SMATrust. CR is supported by a Career Development Fellowship from the Wellcome Trust (205162/Z/16/Z). Work in the laboratory of MJG was supported by the Medical Research Council (MRC programme number U105178803). JRC is supported by a Wellcome Innovator Award (grant number 210774/Z/18/Z). JEM was supported by Great Ormond Street Hospital Children’s Charity (grant number V2137). The support of MDUK and the MRC Centre for Neuromuscular Diseases is also gratefully acknowledged. JM is supported by the MDUK. JRC and JEM are partly funded by the NIHR Great Ormond Street Hospital Biomedical Research Centre. The views expressed are those of the authors and not necessarily those of the NHS, the NIHR or the Department of Health. Thanks are also due to ERN Euro-NMD (www.ern-euro-nmd.eu) to AF as member and Chair of the Genetic Task.

## Author Contributions

CAB managed, led and wrote the project/manuscript. CAB and AJ conceptualised, curated, analysed and reviewed the project. TLEvW performed experiments and statistical analysis. MB performed experiments and assisted with interpretation of results and guidance. SB and GM assisted with interpretation. JM, LC, KM, AB, WFL, CR, KC and JEM assisted with experiments. JRC, EOD, AFS and MJG provided materials for the study. MSF and AF provided patient RNA for study. RGF and MJAW supervised study.

## Supplementary Information

Table S1. SiRNA and PCR Primer IDs or Sequences

**Figure S1.**
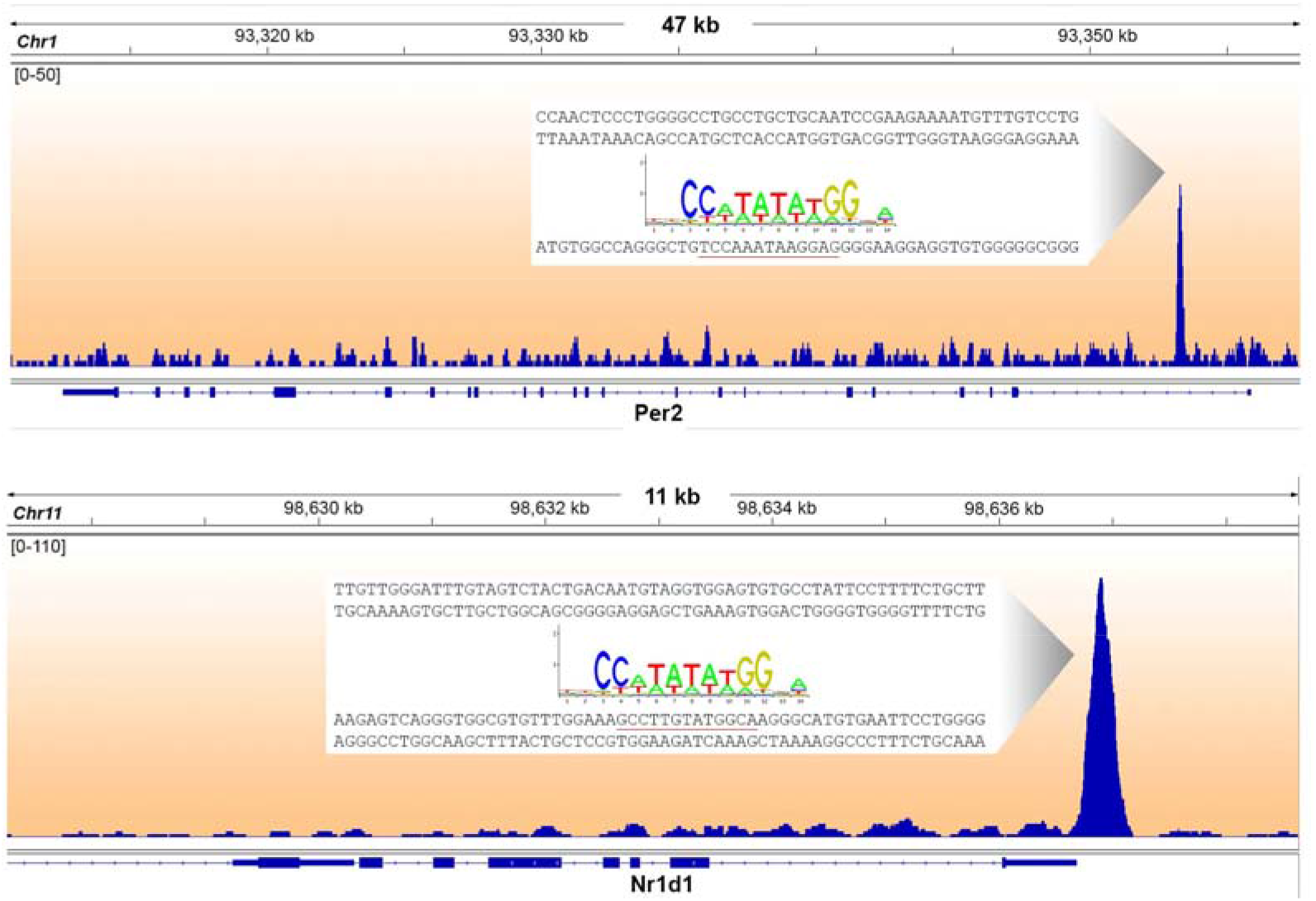
SRF motif binding sites for Per2 and Nr1d1 as determined by Esnault *et al*^9^

**Figure S2.**
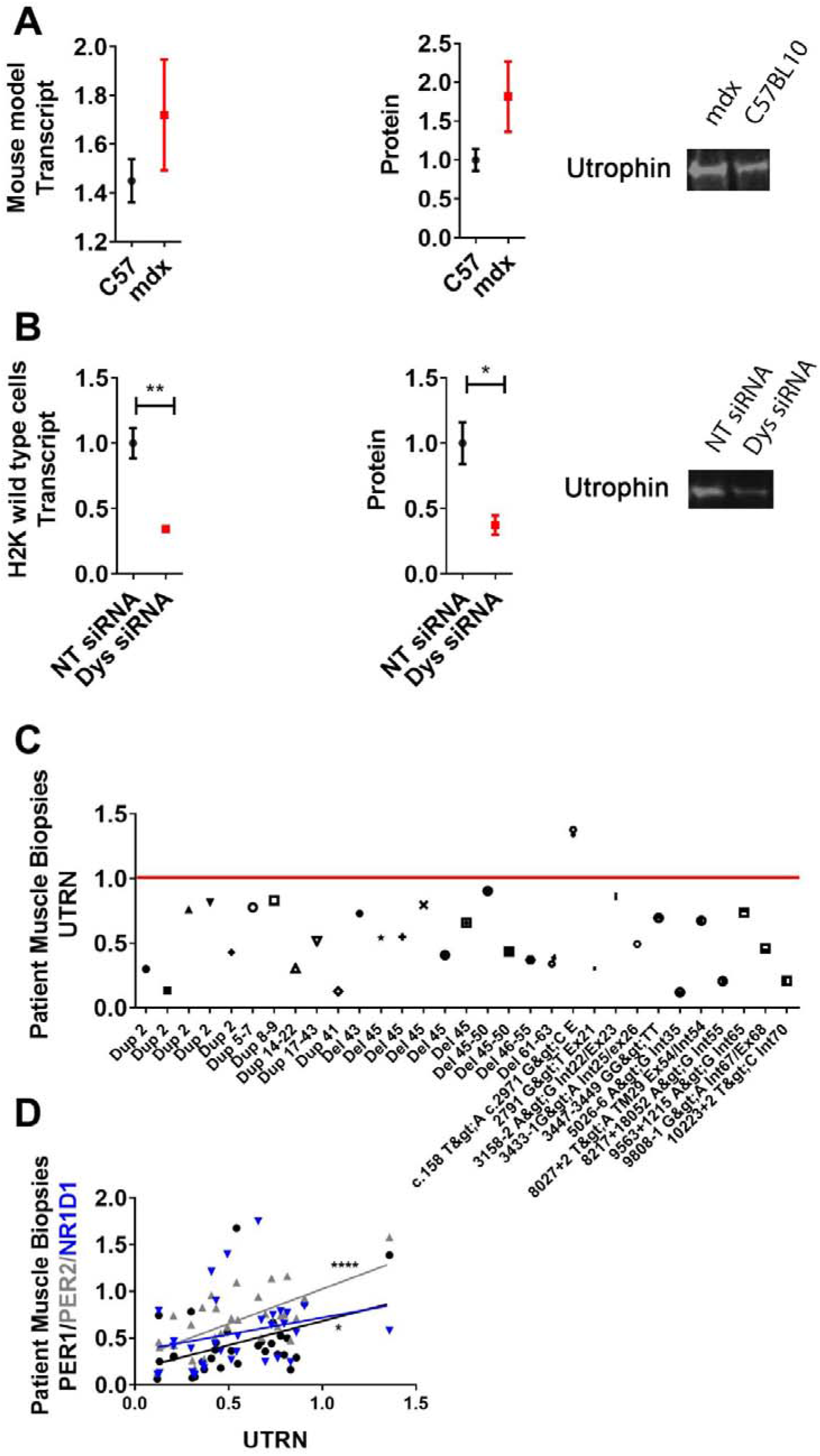
Utrophin protein and transcript levels altered in dystrophic models. Transcript levels (left), western blot quantification (middle) and representative western blots (right; total protein quantified for normalisation) for utrophin shown for **(A)** *mdx* and C57BL10 mice and **(B)** H2K 2B4 motubes. **(C)** *UTRN* gene expression down-regulated in all but one DMD muscle biopsy sample. **(D)** Linear regression plotting *PER1, PER2,* and *NR1D1* versus *UTRN* gene expression indicates significant correlation pattern with *PER1* (R= 0.1687) and *PER2* (R= 0.4542). Data normalised to NT siRNA or C57BL10; Student’s *t*-test for comparisons between 2 groups (Two-tailed; ***p⍰<⍰0.001, **p⍰<⍰0.01, *p⍰<⍰0.05). N=3-4.

**Figure S3.**
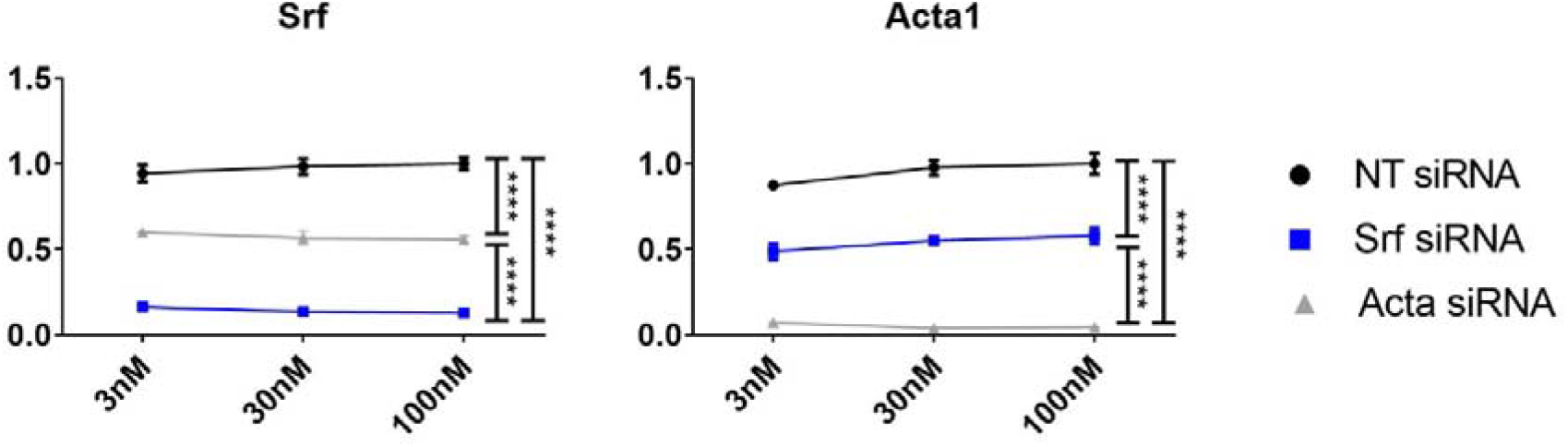
In vitro dose response curves for *Srf* and *Acta*. H2K 2B4 myotubes were transfected with 3nM, 30nM and 100nM of siRNA targeting the *Srf, Acta* and *Dmd* genes (*Srf* siRNA, blue, *Acta* siRNA grey and *Dmd* siRNA, red), and a non-targeting (NT siRNA, black) siRNA was used for control. Gene expression data indicates significant and sustained knock-down of respective genes at low doses (housekeeping gene-*Gapdh*). Data normalised to NT siRNA; n=3; Two-way ANOVA with Bonferroni *post-hoc* test performed. Mean values reported with SEM; ***p⍰<⍰0.001, **p⍰<⍰0.01, *p⍰<⍰0.05.

## References

1. Dunckley, M.G., et al., Modification of splicing in the dystrophin gene in cultured Mdx muscle cells by antisense oligoribonucleotides. Hum Mol Genet, 1998. 7(7): p. 1083–90.

2. Wilton, S.D., et al., Specific removal of the nonsense mutation from the mdx dystrophin mRNA using antisense oligonucleotides. Neuromuscul Disord, 1999. 9(5): p. 330–8.

3. Amann, K.J., A.W. Guo, and J.M. Ervasti, Utrophin lacks the rod domain actin binding activity of dystrophin. J Biol Chem, 1999. 274(50): p. 35375–80.

4. Norwood, F.L., et al., The structure of the N-terminal actin-binding domain of human dystrophin and how mutations in this domain may cause Duchenne or Becker muscular dystrophy. Structure, 2000. 8(5): p. 481–91.

5. Banuelos, S., M. Saraste, and K. Djinovic Carugo, Structural comparisons of calponin homology domains: implications for actin binding. Structure, 1998. 6(11): p. 1419–31.

6. Ohlendieck, K., Towards an understanding of the dystrophin-glycoprotein complex: linkage between the extracellular matrix and the membrane cytoskeleton in muscle fibers. Eur J Cell Biol, 1996. 69(1): p. 1–10.

7. Sproat, B.S., et al., Highly efficient chemical synthesis of 2’-O-methyloligoribonucleotides and tetrabiotinylated derivatives; novel probes that are resistant to degradation by RNA or DNA specific nucleases. Nucleic Acids Res, 1989. 17(9): p. 3373–86.

8. Gerber, A., et al., Blood-borne circadian signal stimulates daily oscillations in actin dynamics and SRF activity. Cell, 2013. 152(3): p. 492–503.

9. Esnault, C., et al., Rho-actin signaling to the MRTF coactivators dominates the immediate transcriptional response to serum in fibroblasts. Genes Dev, 2014. 28(9): p. 943–58.

10. Michele, D.E., F.P. Albayya, and J.M. Metzger, Thin filament protein dynamics in fully differentiated adult cardiac myocytes: toward a model of sarcomere maintenance. J Cell Biol, 1999. 145(7): p. 1483–95.

11. Miller, B.H., et al., Circadian and CLOCK-controlled regulation of the mouse transcriptome and cell proliferation. Proc Natl Acad Sci U S A, 2007. 104(9): p. 3342–7.

12. Gao, H., et al., The clock regulator Bmal1 protects against muscular dystrophy. Exp Cell Res, 2020. 397(1): p. 112348.

13. Lamon, S., M.A. Wallace, and A.P. Russell, The STARS signaling pathway: a key regulator of skeletal muscle function. Pflugers Arch, 2014. 466(9): p. 1659–71.

14. Lahoute, C., et al., Premature aging in skeletal muscle lacking serum response factor. PLoS One, 2008. 3(12): p. e3910.

15. Muses, S., J.E. Morgan, and D.J. Wells, A new extensively characterised conditionally immortal muscle cell-line for investigating therapeutic strategies in muscular dystrophies. PLoS One, 2011. 6(9): p. e24826.

16. Deconinck, A.E., et al., Utrophin-dystrophin-deficient mice as a model for Duchenne muscular dystrophy. Cell, 1997. 90(4): p. 717–27.

17. Zheng, B., et al., The mPer2 gene encodes a functional component of the mammalian circadian clock. Nature, 1999. 400(6740): p. 169–73.

18. Lucassen, E.A., et al., Environmental 24-hr Cycles Are Essential for Health. Curr Biol, 2016. 26(14): p. 1843–53.

19. Iwamoto, A., et al., Effects of chronic jet lag on the central and peripheral circadian clocks in CBA/N mice. Chronobiol Int, 2014. 31(2): p. 189–98.

20. Rybakova, I.N., et al., Utrophin binds laterally along actin filaments and can couple costameric actin with sarcolemma when overexpressed in dystrophin-deficient muscle. Molecular Biology of the Cell, 2002. 13(5): p. 1512–1521.

21. Della Marca, G., et al., Decreased nocturnal movements in patients with facioscapulohumeral muscular dystrophy. J Clin Sleep Med, 2010. 6(3): p. 276–80.

22. Bersanini, C., et al., Nocturnal hypoxaemia and hypercapnia in children with neuromuscular disorders. Eur Respir J, 2012. 39(5): p. 1206–12.

23. Gao, Q.Q. and E.M. McNally, The Dystrophin Complex: Structure, Function, and Implications for Therapy. Comprehensive Physiology, 2015. 5(3): p. 1223–1239.

24. Chahbouni, M., et al., Melatonin treatment normalizes plasma pro-inflammatory cytokines and nitrosative/oxidative stress in patients suffering from Duchenne muscular dystrophy. J Pineal Res, 2010. 48(3): p. 282–9.

25. Guglieri, M., et al., Developing standardized corticosteroid treatment for Duchenne muscular dystrophy. Contemp Clin Trials, 2017. 58: p. 34–39.

26. Tsang, A.H., J.L. Barclay, and H. Oster, Interactions between endocrine and circadian systems. J Mol Endocrinol, 2014. 52(1): p. R1–16.

27. Gnocchi, D. and G. Bruscalupi, Circadian Rhythms and Hormonal Homeostasis: Pathophysiological Implications. Biology (Basel), 2017. 6(1).

28. Morrison-Nozik, A., et al., Glucocorticoids enhance muscle endurance and ameliorate Duchenne muscular dystrophy through a defined metabolic program. Proc Natl Acad Sci U S A, 2015. 112(49): p. E6780–9.

29. Fan, L., et al., Kruppel-like factor 15: Regulator of BCAA metabolism and circadian protein rhythmicity. Pharmacol Res, 2018. 130: p. 123–126.

30. Zhang, L., et al., KLF15 Establishes the Landscape of Diurnal Expression in the Heart. Cell Rep, 2015. 13(11): p. 2368–2375.

31. Jeyaraj, D., et al., Klf15 orchestrates circadian nitrogen homeostasis. Cell Metab, 2012. 15(3): p. 311–23.

32. Walter, L.M., et al., Interventions Targeting Glucocorticoid-Kruppel-like Factor 15-Branched-Chain Amino Acid Signaling Improve Disease Phenotypes in Spinal Muscular Atrophy Mice. EBioMedicine, 2018. 31: p. 226–242.

33. Schroeder, A.M. and C.S. Colwell, How to fix a broken clock. Trends Pharmacol Sci, 2013. 34(11): p. 605–19.

34. Kadener, S., et al., Regulation of alternative splicing by a transcriptional enhancer through RNA pol II elongation. Proc Natl Acad Sci U S A, 2002. 99(12): p. 8185–90.

35. West, S., N. Gromak, and N.J. Proudfoot, Human 5’ --> 3’ exonuclease Xrn2 promotes transcription termination at co-transcriptional cleavage sites. Nature, 2004. 432(7016): p. 522–5.

36. Betts, C.A., et al., Implications for Cardiac Function Following Rescue of the Dystrophic Diaphragm in a Mouse Model of Duchenne Muscular Dystrophy. Sci Rep, 2015. 5: p. 11632.

